# Predicting Rhizosphere Competence Related Catabolic Gene Clusters in plant-associated bacteria with RhizoSMASH

**DOI:** 10.1101/2025.03.29.646099

**Authors:** Yuze Li, Mingxue Sun, Jos M. Raaijmakers, Liesje Mommer, Fusuo Zhang, Chunxu Song, Marnix H. Medema

## Abstract

Plants release a substantial fraction of their photosynthesized carbon into the rhizosphere as root exudates, a mix of chemically diverse compounds that drive microbiome assembly. Deciphering how plants modulate the composition and activities of rhizosphere microbiota through root exudates is challenging, as no dedicated computational methods exist to systematically identify microbial root exudate catabolic pathways. Here we used and integrated published information on catabolic genes in bacterial taxa that contribute to their rhizosphere competence. We developed the RhizoSMASH algorithm for genome-synteny-based annotation of rhizosphere-competence-related catabolic gene clusters (rCGCs) in bacteria by means of a set of 58 knowledge-based logic detection rules carefully curated through sequence similarity network analysis. Our analysis revealed large heterogeneity of rCGC prevalence both across and within plant-associated bacterial taxa, indicating extensive niche specialization. Furthermore, we validated that the presence or absence of rCGCs in bacterial genomes reflects their catabolic capacity and is predictive for their rhizosphere competence by aligning rhizoSMASH results with paired genome/metabolome datasets of rhizobacterial taxa. RhizoSMASH provides an extensible framework for studying rhizosphere bacterial catabolism, allowing targeted selection of beneficial bacterial taxa for microbiome-assisted breeding approaches for sustainable agriculture.

## Introduction

The rhizosphere, the narrow soil zone surrounding and influenced by plant roots, exhibits physicochemical properties distinct from bulk soil, shaped by root-mediated carbon deposition and metabolic activity. Plants release a significant amount of photosynthesized carbohydrates into their rhizosphere^1,2^. These exudates comprise a complex blend of chemical compounds, including sugars, organic acids, amino acids, biogenic amines, and secondary metabolites which can vary substantially across plant species, genotypes within a plant species, developmental stages, and physiological conditions^3–5^. Acting as nutrients, chemoattractants, and antimicrobials, root exudates shape the composition and activities of microbial communities in the rhizosphere^4,6,7^. Bacteria adapted at colonizing this niche, collectively termed rhizobacteria, form part of the plant beneficial microbiome and can enhance nutrient acquisition, regulate phytohormone homeostasis, and mitigate (a)biotic stresses^8–18^.

Given the vast and dynamic chemical diversity of root exudates, deciphering how plants metabolically modulate rhizosphere community composition and activity remains a major challenge^19^. Understanding rhizobacterial catabolic pathways is crucial for predicting host selection of their microbiota and the subsequent rhizobacterial colonization success^20–24^. Some studies have shown that the dysfunction of a single catabolic pathway can directly affect rhizosphere competence, i.e. the ability of microbial taxa to colonize the rhizosphere in competition with other microorganisms^25–28^. Thus, linking bacterial catabolic capacities to plant root exudate profiles is instrumental for predicting rhizosphere competence.

Predicting bacterial catabolic capacities from genomic data remains challenging due to functional diversification within enzyme families. Homology-based approaches often fail to accurately annotate enzyme-coding genes, as evolutionarily divergent enzymes within shared protein families frequently exhibit high sequence similarity yet catalyze distinct reactions. For example, in *Pseudomonas* spp. genomic annotations, genes encoding L-lysine mono-oxygenases tend to be misannotated as L-tryptophan monooxygenase, because they are from the same protein family (PF01593) and have high mutual sequence similarity^15,29,30^. The annotation accuracy of metabolic genes can be improved by leveraging their genomic synteny context^31^. Genes encoding enzymes from a catabolic pathway often co-localize in the genome, forming operons or larger gene clusters. This arrangement not only facilitates coordinated gene expression in polycistronic mRNA(s), but also enables related genes being translocated together as a whole mobile unit^32,33^. This phenomenon enables to distinguishing the above-mentioned L-lysine and L-tryptophan-monooxygenase-coding genes based on whether they cluster with a carbon-nitrogen hydrolase family (PF00795) or an amidase family (PF01425) gene (Patten, Blakney and Coulson, 2013). The ability to accurately and systematically identify rhizosphere catabolic gene clusters (rCGCs) would facilitate establishing functional links between plant metabolic diversity and their microbiome composition and thus enable breeding strategies to steer plant microbiome composition through specific constituents in the root or shoot exudates.

Here, we leverage synteny-based annotation principles to address this challenge, and introduce a new bioinformatic tool, named rhizoSMASH (**rhizo**sphere-competence-related cataboli**SM A**nalysis **SH**ell), which applies a rule-based gene cluster detection algorithm based on the successful principles developed in antiSMASH^34^ to facilitate the prediction of gene clusters related to the catabolism of root exudate metabolites. The first distribution of rhizoSMASH contains over 50 rCGC detection rules, carefully manually curated based on sequence similarity network analysis and covering a wide range of gene clusters encoding pathways that catabolize specific carbohydrates, organic acids, amino acids, biogenic amines, phytohormones, and aromatic compounds found in root exudates. We screened soil- and rhizobacterial genomes with rhizoSMASH and revealed patterns in taxonomic and genomic distribution of rCGCs. Our analyses on two case studies also verified the connection between catabolic capacities and the genomic presence of rCGCs, indicating that the predictive value of rhizoSMASH-based rCGC absence/presence profiles on rhizosphere competence is ‘on par’ with that of experimentally measured substrate utilization assays.

## Results and Discussions

### RhizoSMASH accurately identifies diverse classes of root-associated catabolic gene clusters

We introduced a first working version of the **rhizoSMASH** algorithm that predicts rCGC in bacterial genomes. As a member of the antiSMASH software family^34,35^, rhizoSMASH also predicts gene clusters using a set of detection rules. Each of these rules describes the combination(s) of functional domains captured by profile hidden Markov models (**Figure 1A**, for more details see **Materials and Methods**).

**Figure 1.**
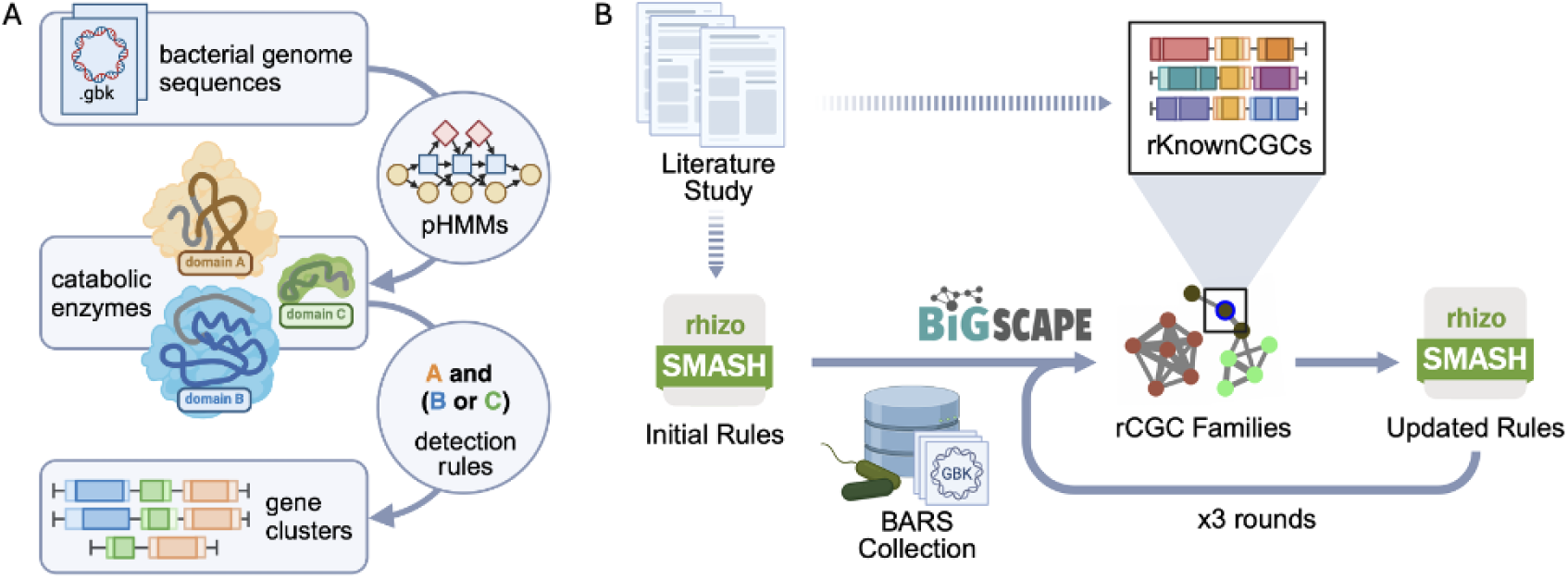
**A.** The gene cluster prediction workflow of rhizoSMASH. RhizoSMASH takes a genome sequence file as input (GenBank or FASTA) and recognize potential catabolic enzymes by scanning the sequence profile hidden Markov models. Gene cluster encoding relevant pathways were then detected using a set of detection rules. **B.** The tuning procedure used for curation of rCGC detection rules. An initial set of detection rules was first summarized from a comprehensive literature study. Then, genome sequences in our BARS collection were scanned using this set of detection rules. The output gene clusters were grouped into cluster families with BiG-SCAPE together with our known cluster database, rKnownCGCs. We manually curated the detection rules by visually investigating the gene cluster family network generated by BiG-SCAPE for putative false positives/negatives, aided by further literature searches when needed. This calibration, validation and finetuning was performed three times to arrive at more and more optimal detection rules.

The development of rhizoSMASH started with an initial set of detection rules summarized from known rCGCs with genetic, and/or biochemical evidence for their function in literature (**Supp.** Figure 1, **Supp. Table 2**). The detection rules then underwent three rounds of manual calibration using sequence similarity network analysis to improve their prediction accuracies based on detailed literature-guided assessment of detected gene cluster families to identify putative false positives and false negatives (**Figure 1B**, final verification results are available at https://www.bioinformatics.nl/~li286/rhizosmash-demo, and more details are in **Materials and Methods**)^36^. The final set consists of 58 detection rules, covering pathways that catabolize six chemical classes of substrates commonly found in root exudates, which are carbohydrates, organic acids, amino acids, amines, phytohormones and aromatic compounds. The explanations of these substrates, their catabolism in bacteria, examples of known rCGCs, and evidence for their relevance in rhizosphere competence are documented at https://www.bioinformatics.nl/~li286/rhizosmash-doc.

In the working version of rhizoSMASH, we largely focused on catabolic pathways of soluble primary metabolites (with several exceptions). However, macromolecules, plant-derived polysaccharides, volatile compounds, and plant-species-specific secondary metabolites have also been shown to affect rhizosphere bacteria-plant interactions^2,7,37–41^. RhizoSMASH is an extendable software similar to the other members from the antiSMASH family, and more detection rule covering catabolism pathways utilizing the above-mentioned metabolite repertoires will be added into rhizoSMASH in the future versions.

### High diversity of catabolic gene clusters across rhizosphere microbiota indicates dynamic metabolic niche adaptation

We constructed a collection of 1,226 genome assemblies from **BA**cteria in the **R**hizosphere and **S**oil (the **BARS** collection, see details in **Materials and Methods** and **Supp. Table 1**) to study the distribution of rCGCs across bacterial taxonomy. We summarized the prevalence of each rCGC type in 20 bacterial families with at least 8 complete assembled genomes in the BARS collection (**Figure 2A**). Our results showed that several rCGC types have high prevalence across almost all bacterial families (e.g. glutamate synthase *glt* cluster, 79.3% and glutamine synthetase *gln* cluster, 93.6%), indicating that these gene clusters may encode pathways of fundamental roles in catabolism that are present in most rhizobacterial genomes. Other rCGC types are specific to a limited range of clades: the L-proline catabolizing *put* clusters are almost only found in *Pseudomonadota* (previously known as *Proteobacteria*) and a few *Bacillota* (previously known as *Firmicutes*) genomes, and the D-proline racemase pathway *prd* clusters are almost limited to *Clostridiaceae*^42^. Some gene cluster types encoding pathways with the same substrate, such as the sucrose hydrolase, phosphorylase, and the levansucrase detour pathways, share overlapping distributions across taxonomy. However, for some other cases, such as the trehalose phosphotransferase and the trehalase pathways, they display complementary distribution, suggesting niche differentiation across different taxa (*phi*=-0.27, Fisher’s exact, *P*=4×10^−7^). Note that the absence of an rCGC in a genome does not necessarily mean the genome does not carry any genes encoding the pathway; in rare cases, these genes may be scattered in the genome and, therefore, not recognized as a gene cluster in rhizoSMASH. Also, because the rCGC detection rules were designed based on known studies, catabolic pathways encoded by undocumented gene cluster may also exist.

**Figure 2.**
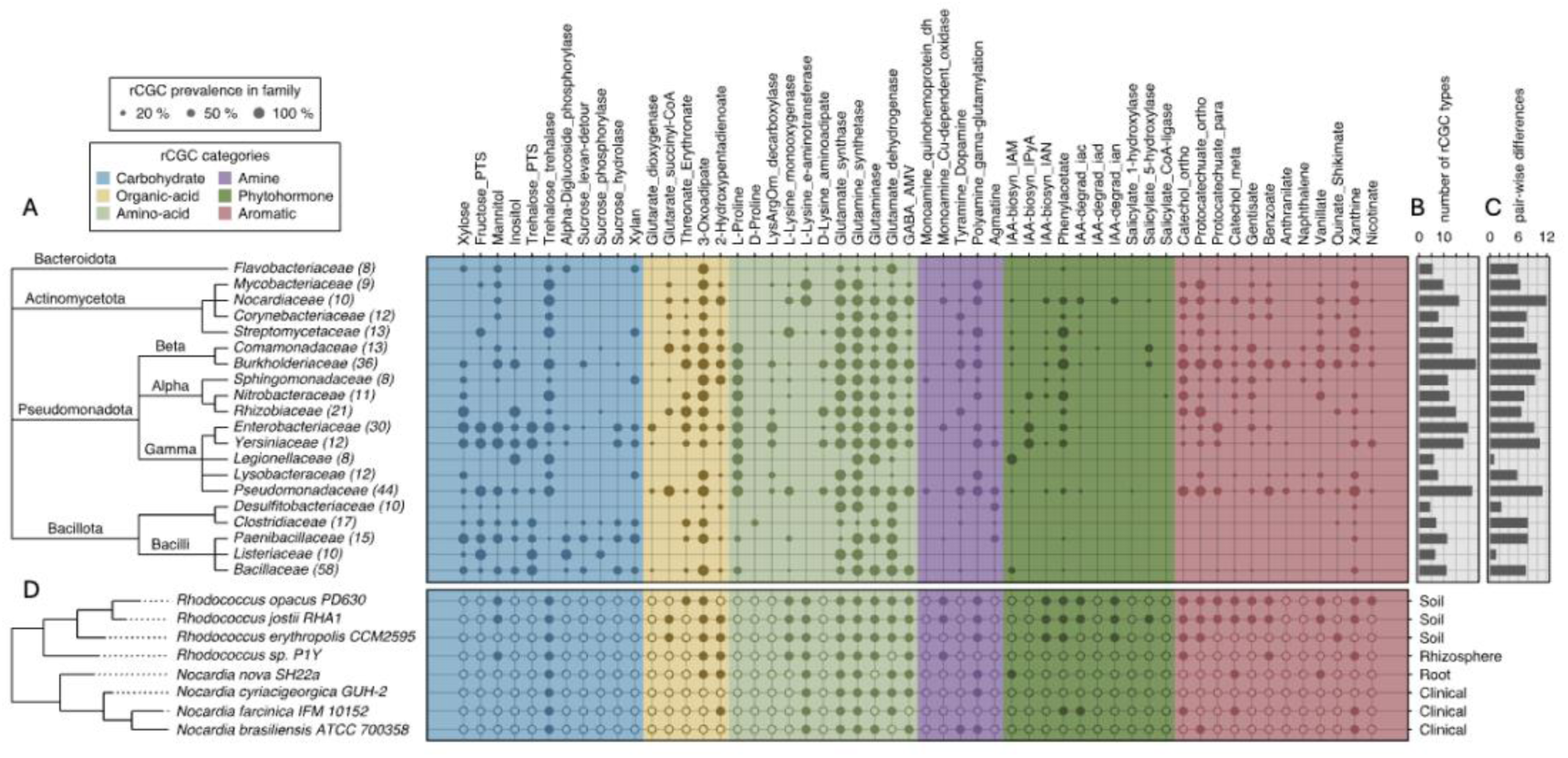
**A.** Distribution of rCGCs across bacterial taxonomy. The size of a dot represents the percentage of genomes in a family that an rCGC type has been detected by rhizoSMASH (defined as prevalence in the main text). The phylogenetic tree on the left was derived from the lineage information available at NCBI. The number in parentheses indicates the number of completely assembled genome in BARS for each family. **B.** The abundance (the average number of rCGC types present in the genomes of a family) and **C.** the diversity (the average differences in rCGC presence/absence between each pair of genomes in a family) indexes of rCGC types across bacterial families. **D.** The rCGC presence/absence profiles in *Nocardiaceae* spp. genomes whose isolation origin were clear, where a solid dot represents the presence of a rCGC type in a genome, and an open dot represents the absence. The labels on the right represent the isolation origins, based on their records at JGI GOLD and NCBI. The phylogeny on the left was constructed with genomic 16S rDNA sequences, rooted with the *Psedomonas putida* type strain NBRC 14164 as an outgroup (not shown on the tree).

Non-metric multidimensional scaling (NMDS) of the prevalence of the rCGC of 20 bacterial families (**Supp.** Figure 2) showed taxonomically related families to share similar rCGC repertoires (PERMANOVA of the null: familywise rCGC prevalences are the same across all phyla, *P*=0.001). Specifically, *Bacillota* genomes contain more carbohydrate-catabolism-associated rCGCs (Wilcoxon, *P*=0.0219), consistent with a previous phylogenetic study on the distribution of carbohydrate-activate enzymes (CAZymes) in bacteria^43^. In contrast, aromatic-compound- and phytohormone-catabolizing rCGCs are more frequently found in *Pseudomonadota* and *Actinomycetota* (Wilcoxon, P<2.2×10^−16^). Indeed, most known bacterial aromatic catabolic pathways^44,45^, as well as auxin biosynthesis and degradation pathways^46,47^, were characterized within these two phyla.

Subsequently, zoomed from a broader taxonomic rank into each bacterial family. To this end, we investigated the abundance and diversity (detailed definitions in Materials and Methods: Family-wise distribution of rCGCs) of rCGC types in each family (**Figure 2B, 2C**). The results showed that families with the highest abundance of rCGCs generally belong to *Pseudomonadota*. Within *Pseudomonadota*, the families *Burkholderiaceae* and *Pseudomonadaceae* have the highest abundance and diversity in rCGC types in their genomes, indicating that many of these organisms are metabolic generalists able to grow on a wide array of root exudate components. These two families are also frequently found as members of the most dominant bacteria families in various rhizosphere microbiomes^48,49^. Especially, the family *Pseudomonadaceae* harbors a large number of known PGPR strains^9,50–56^. In contrast, genomes in the families *Desulfitobacteriaceae*, *Listeriaceae* and *Legionellaceae* displayed both the lowest abundance and diversity in rCGC types (**Figure 2B, 2C**). These families are generally not enriched in the rhizosphere: *Desulfitobacteriaceae* spp. bacteria are strict anaerobic sulfate or organohalide reducers commonly found in anoxic sediment^57,58^; *Listeriaceae* and *Legionellaceae* are families whose members are known as food-borne or environmental pathogens^59,60^. All genome accessions from these three families belong to the RefSoil subcollection, indicating their presence in various soil samples based on metagenomic reads that were mapped to their 16S rDNA sequences^61^, but a detailed inspection of these strains showed that none of them were originally isolated from plant-associated environments including the rhizosphere (**Supp. Table 3**). These results suggest that rhizosphere-dwelling bacteria carry more abundant and diverse rCGCs than bacteria adapted to ecosystems other than plants.

Subsequent focus on the family *Nocardiaceae*, which showed the highest diversity of rCGC types across all tested families (**Figure 2A, D**) suggests that members of this family may have different catabolic strategies to adapt to their local environments. *Nocardiaceae* strains in our study were originally isolated from various ecosystems, including human patients, soil, rhizosphere and root (**Supp. Table 3**) which was reflected in their phylogeny (**Figure 2D**). In general, environmental strains tend to carry more rCGC types compared to clinical *Nocardiaceae* strains (Wilcoxon, P = 0.018). Soil-derived strains are abundant in aromatic compound degradation pathways (**Figure 2D**), which enable them to utilize lignin-derived materials that are abundant in soil^62,63^. The existence of auxin-metabolizing pathways in *Nocardiaceae* spp. has been reported in several studies^17,64–68^. RhizoSMASH also predicted a group of putative rCGCs encoding both auxin biosynthesis and degradation pathways that were enriched in environmental strains (**Figure 2D**), including the phenylacetate degradation *paa* cluster^68^ and the indole-3-acetate (IAA) to catechol *iac* gene cluster^17^. The aldoxime dehydratase gene clusters (detected by the IAA biosynthesis IAN pathway rule) were also reported to have certain level of substrate specificity for indole-3-aldoxime (IAOx) in other *Rhodococcus* strains^65,66^. This intra-family diversity of rCGCs suggests roles of related catabolism in local adaptation to different environments.

Zooming further into the sub-genomic level, we also observed unevenly distributed localization of rCGCs within various bacterial genomes (**Supp.** Figure 3). They form sub-chromosomal regions that were enriched with rCGCs, including cases where rCGCs encoding pathways down- or upstream of each other (e.g., a benzoic acid catabolic gene cluster and the downstream catechol 2,3-cleavage gene cluster), suggesting they may form genomic islands (GI) which can facilitate rapid niche adaptation through concerted horizontal gene transfer^69^, though we did not have clear statistical evidence supporting the relationship between these rCGC-rich regions and tRNA genes (one of the GI markers, **Supp.** Figure 3). To further study this intra-genomic diversity of rCGC, we focused on the genus *Burkholderia*, which typically possesses a multipartite genome containing three large circular replicons^70,71^. The nomenclature of these replicons varies in different study fields, and in our study, we named the large replicons in each genome “chromosome”, “chromid” and “megaplasmid” according to their sequence length in descending order. We analyzed all the replicons from completely assembled *Burkholderia* genomes in BARS with NMDS using the rCGC presence/absence profile in each sequence (**Figure 3A**). The result showed that the large replicons are grouped together according to their type labels (chromosome, chromid and megaplasmid labels) instead of their source genomes, indicating nonrandom localization of certain rCGC classes within the same replicon types (**Figure 3A**). Permutational analysis of variances also confirmed that rCGC presence/absence profiles are significantly different among chromosome, chromid and megaplasmid (PERMANOVA; *P*=0.001) but not among genome sources (*P*=1.00) or species (*P*= 0.985). Meanwhile, rCGCs with different substrate categories also show significant preferences in one or two replicons (**Figure 3B**), with for example aromatic compound rCGCs being enriched in megaplasmids and chromids compared to chromosomes, consistent with replicon functional biases reported in previous studies^71,72^. As secondary replicons (chromid and megaplasmids) have faster evolutionary rates^73^, the enrichment of rCGCs in chromid and megaplasmids indicates their importance in environment adaptation^70^.

**Figure 3.**
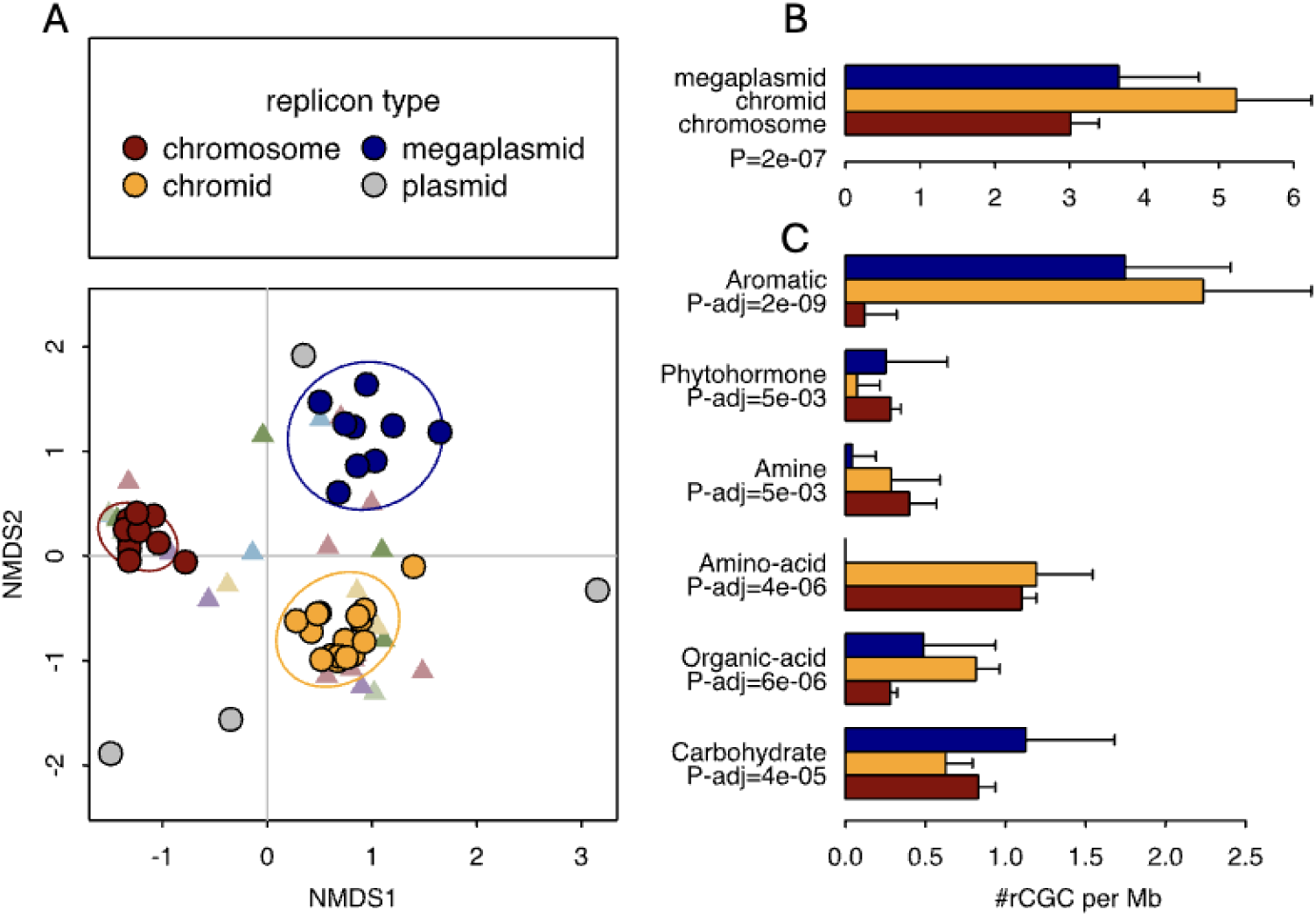
**A.** NMDS plot of *Burkholderia* spp. replicons according to the presence/absence of rCGCs in each replicon sequences. Each point represents a replicon in a strain, while the color of the dot represents the type of replicon (manually labeled according to their sequence lengths). The ellipse represents the 1-SD range for each type of replicon. Each triangle marker represents a rCGC type, colored according to the same schema used in Figure 2. **B.** The average number of rCGCs present per million bases in each replicon type (lengths of error bar = 1-SD). The *P*-value was from the result of a Kruskal-Wallis rank sum test on the equality of three replicon types. **C.** Similar to B., but the rCGC types were divided into six categories based on their substrate properties. The *P*-values were adjusted for multiple testing correction with the Bonferroni method.

### Genomic rCGC profiles predict rhizosphere competence as accurately as experimentally measured catabolism data

Many studies have aimed to predict rhizosphere competence with various approaches and features^4,49,74,75^. Metabolome analyses^4^, whole-genome functional annotations^49,74,75^ or experimental studies with synthetic communities^76,77^ have paved the way to predict rhizosphere behaviors of bacterial groups and have shown clear host-specific connections with specific catabolic activities or genes. However, most of these approaches are quite laborious and do not yet establish direct functional links between root exudate composition and the associated microbial catabolic pathway genes. Here, we use two of these studies^4,20^ to demonstrate the value of the new rhizoSMASH tool for advancing rhizosphere competence analysis based on rCGC profiles.

For our first case study, we adopted the catabolism data for 60 phenazine-producing *Pseudomonas* spp. strains and their rhizosphere colonization data in *Arabidopsis thaliana* and potato rhizospheres published by Zboralski et al. in 2020 (more details in Materials and Methods). The non-metric multidimensional scaling (NMDS) according to both genomic rCGC profiles and assay-measured catabolic capabilities show separation of strains with low rhizosphere competence. These results are in line with our findings above showing the existence of rCGC profile diversity within bacterial families and genus (**Figure 2D**). PERMANOVA also confirmed that both rCGCs profiles and catabolism capabilities were significantly different between high and low (*P*_rCGC_=0.001, *P*_cat_=0.001), and between medium and low (*P*_rCGC_=0.001, *P*_cat_=0.001) rhizosphere colonizers, but not significantly different between high and medium colonizers (*P*_rCGC_=0.354, *P*_cat_=0.061). Therefore, we merged the rhizosphere competence levels “high” and “medium” into one level “med-high” in the subsequent analysis. Then, we trained random forest models to predict colonization levels in both *A. thaliana* and potato rhizospheres. According to an 8-fold nested cross-validation, for rhizosphere colonization in *A. thaliana*, prediction accuracy of the rCGC-based model was 0.884 (SD = 0.106), while the accuracy of the catabolism-assay-based model was 0.848 (SD = 0.115). For rhizosphere colonization in potato, prediction accuracies of the rCGC-based model and the catabolism-assay-based model were 0.783 (SD=0.122) and 0.681 (SD=0.110) (**Figure 4C**). Models trained on rCGC profiles were in general at least as accurate as models trained on experimentally measured catabolism capabilities (**Figure 4C**).

**Figure 4.**
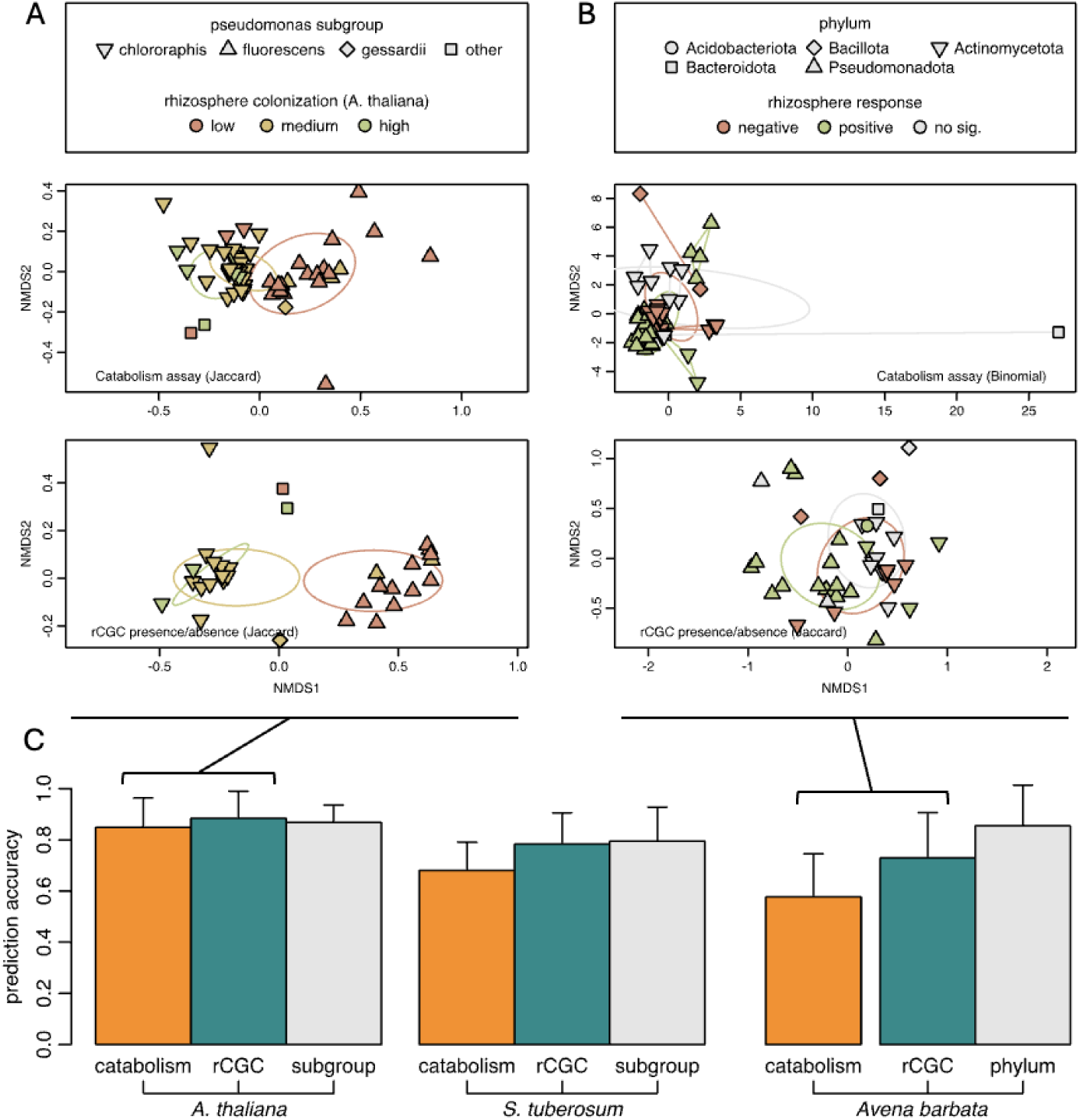
**A.** The NMDS plot of bacterial strains according to their catabolism assay results (top) and genomic rCGC presence/absence profiles (bottom) in the first case study (*Pseudomonas* spp. isolates) and **B.** in the second case study (soil bacterial isolates from Mediterranean grassland). Each point represents a bacterial strain, where the color indicates the rhizosphere competence phenotype, and the shape indicates its A. subgroup in *Pseudomonas* genus and B. Phylum. **C.** The prediction accuracies of prediction models constructed with different datasets, estimated with cross-validations (length of error bar = 1-SD). For the labels on the bottom, the first row indicates the type of the predictors, and the second row indicates the prediction target (rhizosphere competence in the rhizosphere of corresponding plant). Note that the catabolism-assay-based model in the Mediterranean grassland dataset (the orange bar in the right group) was evaluated using a different cross-validation setting, because the catabolism-assay dataset contains repeated observations but on a smaller number of strains; therefore, its accuracy is not comparable within the group.

For our second case study, we then broadened the taxonomic scope of bacteria using the dataset published by Zboralski at al. in 2018, which contains the rhizosphere competence data in *Avena barbata* for a smaller collection of bacterial isolates but from a wider range of phyla compared to the first case study. This dataset also contains root metabolite consumption data in a subset of isolates measured with exometabolomic assays with replications (more details in Materials and Methods). For this more diverse set of strains, the NMDS plots did not show obvious separation of rhizosphere competence levels according to either rCGC profiles or data from catabolism assays (**Figure 4B**), and the PERMANOVA results also reported that no significant differences can be found in the rCGC profiles (*P*_rCGC_=0.128 with Jaccard index) nor for the catabolism assay data (*P*_cat_=1.00 with Binomial index, stratified using bacterial isolate labels) between negative and positive rhizosphere response groups. We also trained random forest models predicting the rhizosphere competence for bacterial isolates with either genomic rCGC profiles or with catabolism assay results (**Figure 4D**). For the rCGC-based model, we achieved an accuracy of 0.729 (SD=0.177), which is lower than the rCGC-based in the *Pseudomonas* limited cases (**Figure 4B, D**). For the catabolism-assay-based model, we only obtained an accuracy of 0.577 (SD=0.169) (**Figure 4D**). However, because the number of bacterial isolates available for training the catabolism-assay-based model was lower compared to those available for the rCGC-based model (explained in Materials and Methods), and different cross-validation methods have been adopted for each of the models, we cannot simply conclude that the catabolism-assay-based one performs worse.

By comparing the rankings of variable importance (**Supp.** Figure 4), we found that for rCGCs with high predictive value in the rCGC-based models, their substrate or related metabolites usually also appear, if measured in the assays, on the list of important variables in the corresponding catabolism-assay base models; these matched pairs include the hemicellulose component xylose, the auxin phenylacetate, the biogenic monoamine phenylethylamine, and the ascorbate degradation downstream organic acid threonate (**Supp.** Figure 4). Within the matched pairs, xylose and phenylacetate are two metabolites that appears on models for all three plant rhizospheres, suggesting their role in rhizosphere colonization may be universal. In contrast, metabolites with plant-specific predictive value, such as L-proline in the second case study, suggest their unique role of mediating colonization in the rhizosphere of that plant. But it should be pointed that high variable importances in random forest models do not always imply a positive correlation, and the size of our case study datasets were also limited to make too general conclusions.

Also, the bacterial strains in these two datasets exhibit a strong phylogenetic bias in rhizosphere competence (**Figure 4A, B**, point shapes). Especially, in the second case study, *Bacillota*, *Pseudomonadota* and *Acidobacteriota* spp. strains were exclusively positive or negative rhizosphere responders. This bias makes a simple taxonomy-based prediction also performed well for both datasets (**Figure 4C**, grey bars). In general, we believe that catabolic gene cluster profiles should not be used as the single source of evidence for predicting rhizosphere competence in general, where other factors such as mobility, biofilm formation, biosynthesis of antibiotics and plant immune response^78–80^ also contribute to determining rhizosphere competence, but the data show that these profiles do contain information with significant predictive value.

After all, competence can be predicted with relatively high accuracy based on a very limited number of binary features without any taxonomic information (**Figure 4C**). Combining rhizoSMASH results with metagenomic and/or metatranscriptomic mapping (facilitated by software such as BiG-MAP software, which can directly take output from any antiSMASH family software as input^81^) in the rhizosphere microbiota when exposed to root exudates with different compositions can be a key next step in unraveling how evolutionary changes in root exudate profiles can lead to shifts in the recruitment of beneficial and non-beneficial microbes by plant hosts, for example during domestication trajectories^82–85^.

### Conclusions

RhizoSMASH is a bioinformatic software package designed to discover catabolic gene clusters in bacterial genomes that are associated with rhizosphere competence. We studied rCGC presence/absence profiles predicted by the working version of rhizoSMASH across a broad range of bacterial taxonomic groups and found functional diversification at various depths of taxonomic ranks (**Figure 2**). We also found uneven local distribution (Supp. Figure 3) and replicon biases (**Figure 3**) of rCGCs throughout genomic sequences in several groups of bacteria. To link rhizoSMASH-detected rCGCs with rhizosphere competence, we performed two case studies, in which we compared the prediction accuracies of rhizosphere competence between rCGC-based and catabolism-assay-based random forest models (**Figure 4**). Results from these two datasets confirm that the presence/absence profiles of the limited collection of rCGCs carry a comparable amount of niche-specific information compared to the catabolism assays that involve a considerably larger number of metabolites. On the whole, these results indicate an important role of these rCGCs in the adaptation to the rhizosphere niche across the phylogenetic tree.

## Materials and Methods

### Construction of the BARS (*Ba*cteria in the *R*hizosphere *S*oil) genome collection

In order to facilitate rhizoSMASH analyses of catabolic diversity across bacterial taxa, we compiled the BARS genome assembly collection, containing genome accessions for soil- and rhizosphere-associated bacteria that are available in the NCBI GenBank database. The version until this manuscript has 1226 genome accessions in total. RhizoBase consists of seven subcollections (REFSOIL, RHIZATHA, SOILATHA, RHIZHVUL, RHIZOSAT, RHIZTAES, RHIZSLYC), with a few overlapping accessions. Among these, the REFSOIL collection contains 842 bacterial entries cited from the RefSoil database^61^; RHIZATHA and SOILATHA came from the at-RSPHERE project which has genome assemblies of 194 Arabidopsis rhizosphere bacteria and 32 bulk soil bacteria^86^; RHIZHVUL, RHIZOSAT, RHIZTAES and RHIZSLYC have 46, 45, 48 and 21 genome accessions respectively, covering bacteria isolated from barley, rice, wheat and tomato rhizosphere. Detailed references and collection sizes are listed in **Supp. Table 1**. These genome accession collections and assembly files (GenBank and FASTA files) can be downloaded and locally reconstructed using scripts at https://git.wur.nl/rhizosmash/rhizosmash-case-studies/-/tree/main/RhizoBase.

### rKnownCGC

The rKnownCGC is a database of annotated known rCGCs that are used as reference gene clusters in the KnownClusterBLAST module in rhizoSMASH. The database was manually compiled: known rCGCs were first summarized in YAML formatted text files, which contain information including the genome accession, genomic position, gene function annotations; then a script was used to automatically download source genome sequences, clip the gene cluster region and add extra annotations to the records. A database file for KnownClusterBLAST can be downloaded using the download-rhizosmash-databases command during the installation of rhizoSMASH. Alternatively, a raw database can be reconstructed using scripts at https://git.wur.nl/rhizosmash/rhizosmash-dev/-/tree/main/rKnownCGC. The version used in the analysis of this manuscript contains 92 hand-curated known rCGC sequences and their annotations, comprising 395 protein-coding genes from 56 GenBank records.

### Workflow of rhizoSMASH

RhizoSMASH adopted the biosynthetic gene cluster detection algorithm from antiSMASH version 6^34^. Similar to other antiSMASH variants such as gutSMASH^35^, rhizoSMASH takes a bacterial genome sequence as input. If the sequence is not annotated, rhizoSMASH will use prodigal to determine the position of gene open reading frames^87^. A curated set of profile hidden Markov models (pHMMs) is used to screen the genome for functional domains in catabolic pathway enzymes^88^. Then a set of “detection rules”, which are logical combinations of pHMMs, is used to identify genome regions that potentially contain any rCGC. Detection rules are divided into 6 categories based on the properties of each pathway’s substrate(s), which are carbohydrate, amino acid, organic acid, amine, phytohormone and aromatic compound. By default, rhizoSMASH knits the outcome into an HTML page that the user can interact with and visually investigate. In the version until this manuscript, the user can activate the KnownClusterBLAST module to identify close matches of rhizoSMASH-identified gene clusters to known rCGCs in the rKnownCGC database. The source code and installation guide for rhizoSMASH can be found at https://git.wur.nl/rhizosmash/rhizosmash.

### Creating and Tuning rCGC detection rules

To create detection rules for rCGCs, we underwent three rounds of rule calibration and curation. We first constructed an initial set of detection rules based on known rCGCs in a preliminary literature search. This set of rules were installed into rhizoSMASH and used to screen RhizoBase genomes. The predicted gene clusters were collected from the output and sent to BiG-SCAPE (a customized version 1 was used the first round; version 2 was used in the second and third rounds) together with known rCGCs. BiG-SCAPE groups gene clusters into cluster families based on their mutual similarities and generates interactive sequence similarity networks. The outputs from BiG-SCAPE were then manually analyzed and verified; we visually investigated if the detected gene clusters were grouped with any known clusters or if they had similar genomic context with any known clusters. The detection rules were then modified to rule out false positive detections and avoid overlap between rules. The updated rules were re-installed into rhizoSMASH and underwent another round of verification repeatedly.

The final round of verification used a deduplicated subset of the BARS collections. The mutual similarity of genome sequences in BARS was estimated using mash^89^; BARS genomes were first indexed using mash sketch with additional options -k 32 (k-mer size 32); their mutual similarities were then estimated using mash dist. BARS genomes showing mash similarity indexes less than 0.15% were merged, where the most completely assembled genome was kept as representative. Details can be found in the documentation of this repository at https://git.wur.nl/rhizosmash/rhizosmash-case-studies/-/tree/main/gene-cluster-distribution.

### Family-wise distribution of rCGCs

To summarize the taxonomic distribution of rCGCs across BARS genomes, we first selected BARS entries for bacterial families that have sufficient number (≥ 8) of completely assembled genomes. An rCGC presence/absence table of these genomes in these families was obtained from the rhizoSMASH output in the final rule verification round. Based on the presence/absence table, we calculated in each family the **prevalence** of each rCGC type, the **abundance** of all rCGC types, and the **diversity** of rCGCs between genomes. The **prevalence** of an rCGC in a family is defined as the percentage of genomes where the rCGC type is found (**Figure 2A**). The rCGC **abundance** of a family is represented by the average count of rCGC types present per genome in the family **(Figure 2B**). The **diversity** of rCGCs in a family is reflected by the average differences in the presence/absence profiles between each pair of genomes in the family.

### Phylogeny trees

The phylogenic tree for bacterial families were simply built based on lineage information obtained from NCBI with a script using the python module ete3. The lineage information was loaded into R and converted into a data.frame. And the tree was made by applying the as.phylo function in the ape package on that data.frame. The phylogenic tree for *Nocardiaceae* spp. strains was built according to their 16S rDNA sequences. We first detected and extracted 16S rDNA from the genomic DNA sequences of eight *Nocardiaceae* spp. strains (combined with an outgroup strain *Pseudomonas putida* NBRC 14164) with barrnap^90^. Multiple sequence alignment of these 16S rDNA sequences was performed with MUSCLE^91^. We used FastTree to generate a phylogeny tree from the multiple sequence alignment^92^. The phylogeny tree was loaded into R and rooted with the root function in the ape package with the parameter resolve.root set to TRUE. The outgroup tip was removed before final visualization.

### Source and properties of datasets used in case studies

The dataset of 60 phenazine-producing *Pseudomonas* isolates used in case study 1 was published by Zboralski at al. in 2018. All 60 isolates in this dataset have complete genome assemblies available in GenBank. Their catabolic capacity data were measured using BIOLOG phenotype microarrays. Their rhizosphere colonization strength data were measured in gnotobiotic systems for two plant species (*Arabidopsis thaliana* and potato) using quantitative PCR targeting DNA from a conserved phenazine biosynthesis gene. The results were categorized into “high”, “medium” and “low” levels.

The Mediterranean grassland soil bacteria dataset used in case study 2 was published by Zhalnina et al. in 2018. This dataset has 39 sequenced isolates with assembly completeness varying from contig to scaffold levels. Their genome assemblies are available on JGI genome portal under GOLD study ID Gs0017561 or proposal ID 653. The catabolic capacity data in this dataset was obtained with an exometabolomic method: they harvested *Avena barbata* root exudates and compared the percent change of each component before and after bacteria cultivation in the exudates with mass spectrometry. Rhizosphere colonization levels in this dataset were quantified based on enrichment of rhizosphere 16S rDNA in response to the growth of *Avena fatua* during a time course from 0 to 12 weeks. Twenty-seven out of 39 isolates in this dataset showed a significantly positive (n=19) or negative (n=8) rhizosphere response, within which, 12 isolates have catabolism data (2 to 4 replications for each isolate).

### Prediction Models

We used R package randomForest to train prediction models for rhizosphere competence. The caret and rsample packages were used for facilitating cross-validation. In case study 1, we merged “high” and “medium” rhizosphere colonization levels into “med-high” to solve imbalance in the data labeling. In-total, 4 random forests were trained with either rCGC presence/absence data or catabolism data to predict rhizosphere competence levels in *Arabidopsis thaliana* or potato. An 8-fold cross validation was performed for each model to optimize the hyperparameter mtry within a grid. The out-of-sample prediction accuracies for these models were further estimated using an outer layer of 8-fold nested cross-validation. Similar settings were adopted in case study 2 but with different methods for cross-validation due to a smaller data size and different approaches were applied in the rCGC-based and the catabolism-assay-based models. For the rCGC-based model, the hyperparameter tuning was done with the built-in out-of-bag (OOB) method and a nested 8-fold cross-validation was applied to estimate the prediction accuracy. For the catabolism-assay-based model, to prevent information leakage due to repeated catabolism measurements, the hyperparameter tuning was done by repeating a 2-fold grouped cross-validation (GCV) 4 times, and the prediction accuracy was estimated by repeating a 4-fold GCV 2 times. Data partition for the regular cross-validations was done with the createFolds function in the caret package. For the GCVs, it was done with the group_vfold_cv function in the rsample package, where the strain names and rhizosphere competence labels were set to the group and strata as parameters

We also used a simple taxonomy-based method to predict rhizosphere competence phenotypes. These predictions were made by finding the dominant phenotype in each taxonomy label. We applied 8-fold cross-validation to estimate the accuracy of this method, where in each training fold, the dominant phenotype in each taxonomy label was recalculated.

### Statistical tests

We used Fisher’s exact test to test the complimentary distribution of two trehalose rCGCs across BARS genomes (**Figure 2A**). We used the one-sided Wilcoxon Rank Sum test to compare the genomic abundance of rCGC types (number of present rCGC types in genomes) between bacterial phyla (**Figure 2A**) and environmental and clinical *Nocardiaceae* spp. genomes (**Figure 2B**), and the Kruskal-Wallis rank sum test to test if there are differences in the presence of rCGC (count per million bases in genomic sequences) between replicons in *Burkholderia* spp. genomes (**Figure 3B, C**). These statistical tests were performed with the fisher.test, the wilcox.test function and kruskal.test function in the stats package in base R. P-values in the *Burkholderia* replicon study was adjusted with the Bonferroni method with the p.adjust function from the stats package in base R. To test whether rCGCs are evenly distributed on chromosomes (**Supp.** Figure 3), we applied a Monto Carlo simulation of 999 repetitions of the mean nearest neighborhood distance (MNN) from a uniform distribution. The p-values were estimated by the fraction of repetitions where the simulated MNN was smaller than the observed MNN.

We used the metaMDS function from the R package vegan to perform NMDS throughout our study. To study the distribution of rCGC type across bacterial taxonomy, the binomial index was used to measure the dissimilarity of rCGC prevalences between phyla (**Supp.** Figure 2). To study the functional biases in replicons of *Burkholderia*, we first labeled the replicons in each genome according to their the Jaccard index was used to measure the dissimilarity of rCGC presence/absence profiles in the (**Figure 3B**). Similarly, the Jaccard index was used in the NMDS of rCGC profiles (**Figure 4A, B**). For the catabolism-assay-derived data, the Jaccard index was used in the first case study and the binomial index was use in the second case study according to their type of data (**Figure 4A, B**). PERMANOVAs were conducted with the adonis2 function also from the vegan package and the same dissimilarity indexes were used as in the NMDS analyses. Besides, in the PERMANOVA of metabolite consumption in bacterial strains in case study 2, because of repeated measurements, we added strain labels as strata to the parameter to ensure PERMANOVA was performed between strains. The number of permutations was set to 999 as the default value in all the PERMANOVA tests.

