## Supplemental Information for "Predicting Rhizosphere Competence Related Catabolic Gene Clusters in plant-associated bacteria with RhizoSMASH"

| **Collection** | **# Acc** | | **Project Number** | **Ref** |
| --- | --- | --- | --- | --- |
|  | **Tot** | **Sub** |  |  |
| REFSOIL | 842 | 842 | n. a. | ^1^ |
| RHIZATHA | 194 | 194 | PRJNA297942 | ^2^ |
| SOILATHA | 32 | 32 | PRJNA298127 |  |
| RHIZHVUL | 46 | 41 | PRJEB42773 | ^3^ |
|  |  | 1 | PRJNA908138 | ^4^ |
|  |  | 1 | PRJNA507263 | ^5^ |
|  |  | 1 | PRJNA344944 | ^6^ |
|  |  | 2 | PRJNA67537, PRJNA67535 | ^7^ |
| RHIZOSAT | 45 | 20 | PRJNA701950 | ^8^ |
|  |  | 1 | PRJNA905546 | ^9^ |
|  |  | 1 | PRJNA880516 | ^10^ |
|  |  | 1 | PRJNA823515 | ^11^ |
|  |  | 1 | PRJNA821328 | ^12^ |
|  |  | 1 | PRJDB8374 | ^13^ |
|  |  | 1 | PRJNA531785 | ^14^ |
|  |  | 1 | PRJNA494794 | ^15^ |
|  |  | 1 | PRJNA454179 | ^16^ |
|  |  | 1 | PRJDB5588 | ^17^ |
|  |  | 1 | PRJNA414941 | ^18^ |
|  |  | 1 | PRJNA379921 | ^19^ |
|  |  | 1 | PRJNA352539 | ^20^ |
|  |  | 1 | PRJNA263863 | ^21^ |
|  |  | 1 | PRJNA168054 | ^22^ |
|  |  | 7 | PRJNA430105 | n. a. |
|  |  | 4 | PRJNA1047279, PRJNA942581, PRJNA597415, PRJNA544607 |  |
| RHIZTAES | 48 | 11 | PRJNA667745 | ^23^ |
|  |  | 7 | PRJNA745065 | ^24^ |
|  |  | 3 | PRJNA643659 | ^25^ |
|  |  | 3 | PRJNA273703, PRJNA275697, PRJNA275699 | ^26^ |
|  |  | 3 | PRJNA67535, PRJNA67537, PRJNA67539 | ^7^ |
|  |  | 1 | PRJNA245780 | ^27^ |
|  |  | 1 | PRJNA309751 | ^28^ |
|  |  | 1 | PRJNA435479 | ^29^ |
|  |  | 1 | PRJNA488819 | ^30^ |
|  |  | 1 | PRJNA531142 | ^31^ |
|  |  | 1 | PRJNA623691 | ^32^ |
|  |  | 1 | PRJNA635904 | ^33^ |
|  |  | 1 | PRJNA71317 | ^34^ |
|  |  | 1 | PRJNA728132 | ^35^ |
|  |  | 1 | PRJNA741525 | ^36^ |
|  |  | 1 | PRJNA78839 | ^37^ |
|  |  | 1 | PRJNA824280 | ^38^ |
|  |  | 1 | PRJNA886615 | ^39^ |
|  |  | 1 | PRJNA605675 | ^40^ |
|  |  | 7 | PRJNA960949, PRJNA202956, PRJNA545192, PRJNA635401, PRJNA714046, PRJNA721340, PRJNA728357 | n. a. |
| RHIZSLYC | 21 | 3 | PRJEB19151, PRJEB19165, PRJEB19184 | ^41^ |
|  |  | 2 | PRJNA603418, PRJNA604468 | ^42^ |
|  |  | 1 | PRJNA239153 | ^43^ |
|  |  | 1 | PRJNA264437 | ^44^ |
|  |  | 1 | PRJNA780183 | ^45^ |
|  |  | 1 | PRJNA795281 | ^46^ |
|  |  | 1 | PRJNA940804 | ^47^ |
|  |  | 1 | PRJNA509704 | ^48^ |
|  |  | 1 | PRJNA531642 | ^49^ |
|  |  | 2 | PRJNA397822 | n. a. |
|  |  | 7 | PRJNA602778, PRJNA602780, PRJNA264433, PRJNA264436, PRJNA646039, PRJNA695080, PRJNA993421 |  |

**Supp. Table 1** References and collection sizes of subcollections in the BARS genome collection. An n. a. mark in the reference column means the genomes were only documented sunder project numbers.

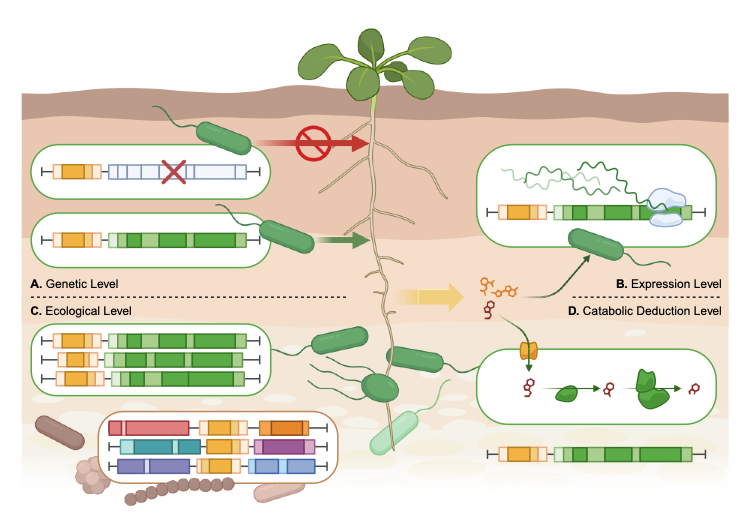

**Supp. Figure 1.** A schematic representation showing the four evidence levels in our literature-based discoveries of known rCGCs. **A.** genetic level: interruption of the gene cluster directly affects rhizosphere competence; **B.** expression level, treatment of root exudates (or root exudate components) induces the expression of the gene cluster; **C.** ecological level, the gene cluster is enriched in the genomes of rhizosphere associated bacteria; **D.** catabolic deduction level, the gene cluster encodes a pathway to catabolize metabolites that are highly abundant in root exudates or are often utilized by rhizosphere associated bacteria.

| Substrate Category | rCGC name | A | B | C | D |
| --- | --- | --- | --- | --- | --- |
| Carbohydrate | Xylose | ^50^ | ^50–52^ | ^50,53,54^ | ^50,51,54^ |
|  | Fructose PTS |  | ^51,52^ |  | ^51^ |
|  | Mannitol |  | ^51,55,56^ | ^50^ | ^51,57^ |
|  | Inositol | ^58,59^ | ^51,55,56,58,60^ | ^58,61^ | ^51,62^ |
|  | Trehalose PTS |  |  | ^54^ | ^51,54,63^ |
|  | Trehalose trehalase | ^59,64^ |  |  |  |
|  | Alpha-Diglucoside phosphorylase |  |  |  | ^51,63^ |
|  | Sucrose levan-detour | ^65^ |  | ^54^ | ^51,54,63,65^ |
|  | Sucrose phosphorylase | ^59^ |  |  |  |
|  | Sucrose hydrolase |  |  |  |  |
|  | Xylan |  | ^55^ |  | ^50^ |
| Organic acid | Glutarate dioxygenase |  |  |  | ^66,67^ |
|  | Glutarate succinyl-CoA |  |  |  |  |
|  | Threonate Erythronate |  |  |  | ^66,68^ |
|  | 3-Oxoadipate | ^69^ | ^51^ |  | * |
|  | 2-Hydroxypentadienoate |  | ^51^ |  |  |
| Amino acid | L-Proline |  | ^70^ |  | ^51,63^ |
|  | D-Proline |  |  |  | ^63,71^ |
|  | LysArgOrn decarboxylase |  | ^56^ |  | ^51,63,68,72^ |
|  | L-Lysine monooxygenase |  |  |  |  |
|  | L-Lysine e-aminotransferase |  |  |  |  |
|  | D-Lysine aminoadipate |  |  |  | ^54^ |
|  | Glutamine synthetase |  | ^55^ |  | ^51,63,73^ |
|  | Glutamate synthase |  | ^51,55,74^ |  |  |
|  | Glutaminae |  |  |  |  |
|  | Glutamate dehydrogenase |  |  |  |  |
|  | GABA/AMV | ^75^ | ^51,75^ |  | ^51,66^ |
| Amine | Monoamine quinohemoprotein dh |  |  |  | ^54^ |
|  | Monoamine Cu-dependent oxidase |  |  | ^54^ |  |
|  | Tyramine/Dopamine |  |  | ^54^ | ^54^ |
|  | Polyamine gama-glutamylation | ^76,77^ | ^51^ | ^54^ | ^54^ |
|  | Agmatine |  | ^51^ |  |  |
| Phytohormone | IAA-biosyn IAM | ^78,79^ |  | ^54,80^ | ^4,51,81–83^ |
|  | IAA-biosyn IPyA | ^84^ | ^85,86^ | ^54,80^ |  |
|  | IAA-biosyn IAN |  |  | ^54,80^ |  |
|  | Phenylacetate |  | ^51,74^ | ^54,87^ |  |
|  | IAA-degrad iac | ^88^ | ^89^ |  | ^63,66,90^ |
|  | IAA-degrad iad |  | ^91^ | ^91^ |  |
|  | IAA-degrad ian |  |  | ^87^ |  |
|  | Salicylate 1-hydroxylase |  |  |  | ^63,66,92,93^ |
|  | Salicylate 5-hydroxylase | ^94^ | ^94^ | ^87^ |  |
|  | Salicylate CoA-ligase |  |  |  |  |
| Aromatic | Catechol ortho |  | ^51^ |  | **  ^54,63^ |
|  | Protocatechuate ortho | ^69^ | ^51,95^ |  |  |
|  | Protocatechuate para |  | ^51^ |  |  |
|  | Catechol meta |  |  |  |  |
|  | Gentisate |  |  |  |  |
|  | Benzoate | ^69^ | ^51^ | ^87^ |  |
|  | Anthranilate |  | ^51^ |  |  |
|  | Naphthalene |  |  |  | ^96^ |
|  | Vanillate |  |  |  | ^63^ |
|  | Quinate/Shikimate |  |  |  | ^51,63,66^ |
|  | Xanthine |  |  |  | ^63^ |
|  | Nicotinate |  |  |  | ^63,66^ |

**Supp. Table 2.** Substate categories, evidence level and references of rCGC detection rules (strict rules, n=54) in the working version of rhizoSMASH. Gene clusters marked with * and ** are downstream and central pathways for aromatic compound catabolism.

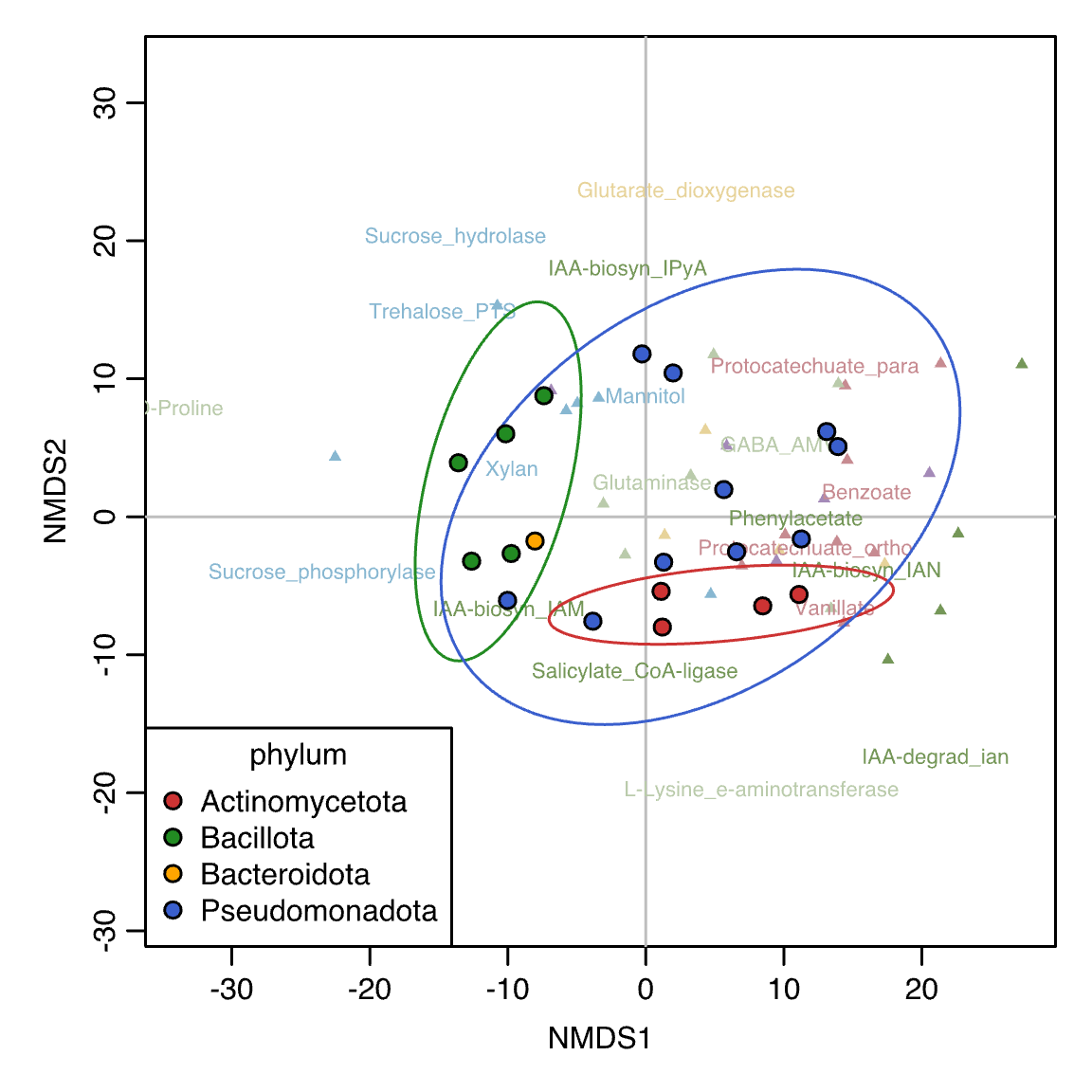

**Supp. Figure 2.** NMDS of bacterial families according to the prevalence of each rCGC type. Each point on the plot stands for a family in Figure 2 in the main text. The color of a point represents its phylum. Phyla of these families are grouped with ellipses, representing the 95% confidence interval of the group. Text labels and triangles represents rCGC types, with the same color scheme as in Figure 2.

| **GenBank Acc.** | **Species** | **Strain** | **Isolated From** |
| --- | --- | --- | --- |
| GCA_000021925.1 | *Desulfitobacterium hafniense* | DCB-2 | Municipal sludge, Denmark |
| GCA_000010045.1 |  | Y51 | Soil contaminated with tetrachloroethene in Japan |
| GCA_000243155.3 | *Desulfitobacterium dehalogenans* | ATCC 51507 | Sediment in freshwater pond, Athens, GA |
| GCA_000243135.3 | *Desulfitobacterium dichloroeliminans* | LMG P-21439 | Soil matrix of an anoxic water-saturated layer (1 m in depth) that had been exclusively polluted with 1,2-DCA |
| GCA_000235605.1 | *Desulfosporosinus orientis* | DSM 765 | Soil at pumping station near rising main Rangoon Road |
| GCA_000255115.3 | *Desulfosporosinus acidiphilus* | SJ4 | Acid mining effluent decantation pond |
| GCA_000231385.3 | *Desulfosporosinus meridiei* | DSM 13257 | Groundwater contaminated with aromatic compounds from motor fuel in sandy soil from Eden Hill, Swan Coastal Plain, Perth, Western Australia |
| GCA_000512895.1 | *Dehalobacter restrictus* | DSM 9455 | PCE-dechlorinating packed-bed column; The Netherlands |
| GCA_000027145.1 | *Listeria seeligeri* | SLCC3954 | Soil, Germany |
| GCA_000008285.1 | *Listeria monocytogenes* | F2365 | Cheese product that caused an outbreak of listeriosis among patients with AIDS in California in 1985 |
| GCA_000168635.2 |  | J0161 | A deli in the United States in 2000 during an epidemic outbreak |
| GCA_000022925.1 |  | 08-5923 | Blood, clinical isolate during a nationwide outbreak |
| GCA_000093125.2 |  | 08-5578 | Blood, reference outbreak strain during a national outbreak |
| GCA_000209755.1 |  | L99 | Cheese; Netherlands |
| GCA_000021185.1 |  | HCC23 | Channel catfish |
| GCA_000218305.1 |  | M7 | Cow’s milk; China |
| GCA_000210815.2 |  | SLCC2372 | spinal fluid of man with cerebrospinal meningitis |
| GCA_000307615.1 |  | SLCC2378 | Poultry |
| GCA_000195795.1 | *Listeria innocua* | Clip11262 | Dairy products (cheese) from Morocco |
| GCA_000210795.2 | *Listeria monocytogenes* | SLCC2482 | Human |
| GCA_000091785.1 | *Legionella longbeachae* | NSW150 | Human |
| GCA_000092545.1 | *Legionella pneumophila* | Corby | Human |
| GCA_000048645.1 |  | Paris | Human, endemic in France |
| GCA_000239175.1 |  | ATCC 43290 | Human lung tissue; USA |
| GCA_000092625.1 |  | 2300/99 Alcoy | Patient affected by legionellosis in Alcoy (Spain) |
| GCA_000048665.1 |  | Lens | Human; France |
| GCA_000250675.3 | *Nocardia brasiliensis* | ATCC 700358 | Human mycetoma, Monterrey, Mexico |
| GCA_000284035.1 | *Nocardia cyriacigeorgica* | GUH-2 | Human nocardiosis case |
| GCA_000009805.1 | *Nocardia farcinica* | IFM 10152 | Bronchus of a 68-year-old male Japanese patient |
| GCA_000523235.1 | *Nocardia nova* | SH22a | Root of *Couma macrocarpa* in Brazil |
| GCA_000454045.1 | *Rhodococcus erythropolis* | CCM2595 | Soil |
| GCA_000014565.1 | *Rhodococcus jostii* | RHA1 | Soil contaminated with gamma-hexachlorocyclohexane in Japan |
| GCA_000599545.1 | *Rhodococcus opacus* | PD630 | Soil sample in Germany |
| GCA_003641205.1 | *Rhodococcus sp.* | P1Y | Rhizosphere of rice; Russia |

**Supp. Table 3.** The environmental origins of bacterial strains from families *Desulfitobacteriaceae*, *Listeriaceae*, *Legionellaceae* and *Nocardiaceae*. We searched the JGI GOLD database with GenBank assembly accession number and full strain names in the corresponding families and summarized “sample collection site”, “isolation country/ocean”, “ecosystem category” and “ecosystem type” fields from the results, combined with their annotations on NCBI. Strains without detailed annotations for these fields were excluded from the table.

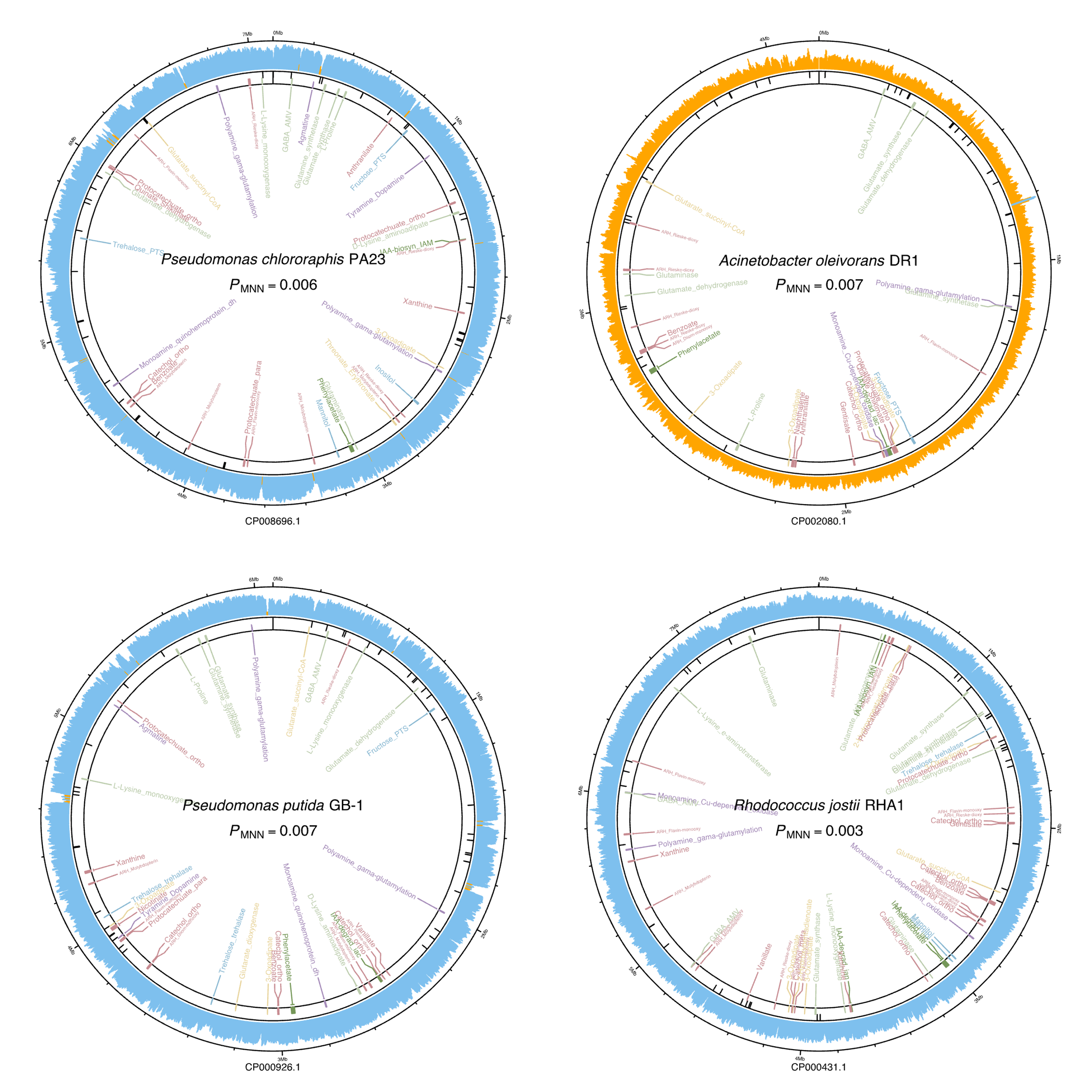

**Supp. Figure 3.** Distribution of rCGCs on the chromosome of four example bacterial strains. The outer ring shows the GC content (with a sliding window of 5 kbps and step of 2 kbps), where blue and orange indicate if local GC content is above 0.5 or not. Black sticks on the middle ring depict the positions of tRNA genes (normally found at the boundaries of genomic islands). The colored sticks on the inner ring indicate the positions of rCGCs, where the colors follow the same scheme in Figure 2. Text labels inside the ring are the name of rCGCs, where smaller text labels are relaxed rules (for the working version, those are core gene clusters encoding the aromatic ring hydroxylating enzymes). Relaxed rCGCs inside the region of other rCGCs are merged into the including rCGC. For each genome, a Monte Carlo simulation of the mean nearest neighborhood distance (MNN) of a uniform distribution was applied to test whether the rCGCs were evenly distributed or not (*P*_MNN_ values under the name of the strain).

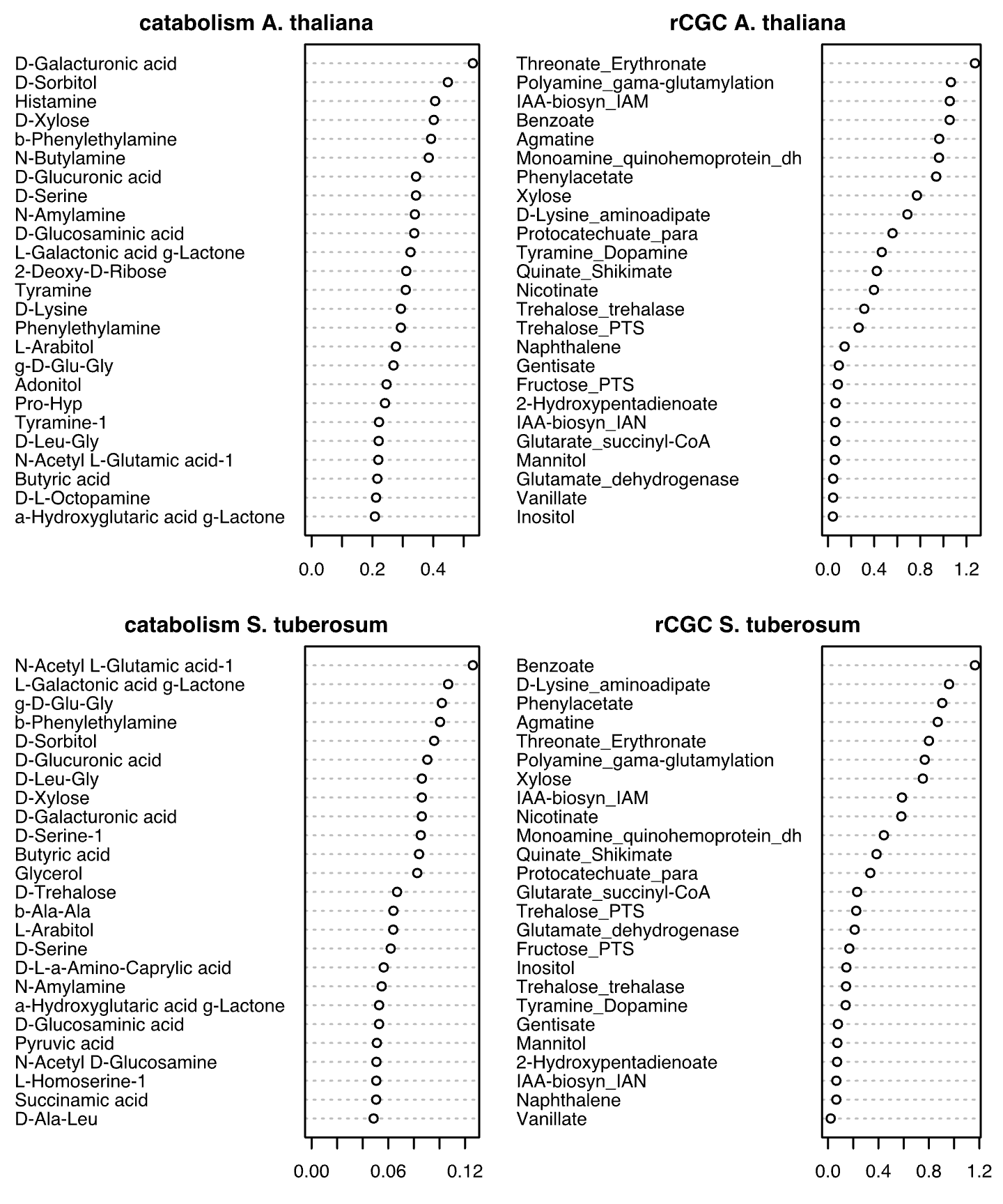

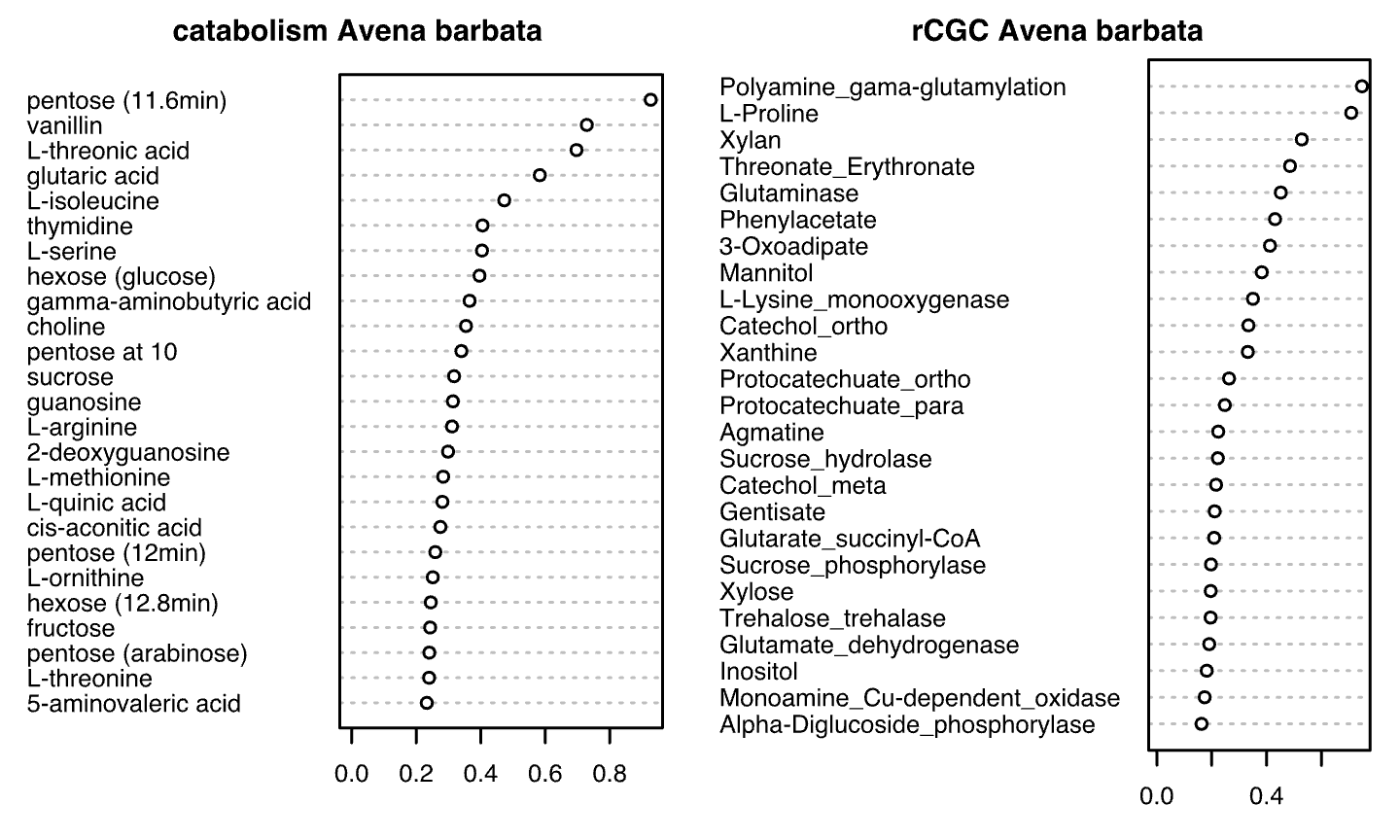

**Supp. Figure 4.** Variable importance charts for random forest models predicting rhizosphere competence phenotypes in different plants and with different datasets. For each model, only the top 25 important predictors are listed on the charts.
